# Detection of immune system activation in hemolymph of *Drosophila* larvae exposed to chitosan-coated magnetite nanoparticles

**DOI:** 10.1101/324335

**Authors:** Doris Vela, Jonathan Rondal, Alexis Debut, Karla Vizuete, Fernanda Pilaquinga

## Abstract

*Drosophila melanogaster* hemolymph cells are confirmed as a model to study the activation of immune system due to foreign stimuli like iron nanoparticles. The toxicity of nanoparticles is a cause for concern due to their effect on human health and the environment. The aim of this study was to detect the activation of cellular immune response in *Drosophila* larvae through the observation of hemolymph composition, DNA damage and larval viability, after the exposure to 500 ppm and 1000 ppm chitosan-coated magnetite nanoparticles for 24 hours. Our results showed activation of cellular immune response after exposure to the nanoparticles owing to the increment of hemocytes, the emergence of lamellocytes and the presence of apoptotic hemocytes. In addition, chitosan-coated magnetite nanoparticles produce DNA damage detected by comet assay as well as low viability of larvae. No DNA damage is showed at 500 ppm. The cellular toxicity is directly associated with 1000 ppm.

## Introduction

*Drosophila melanogaster* has proved to be a suitable organism to test toxic effects of different chemical elements due to its short life cycle and abundant offspring. In *Drosophila* there are two main components of the innate immune response: the humoral and cellular systems, both of which are activated upon immune challenge. The cellular response refers to processes such as phagocytosis, encapsulation, and clotting that are directly mediated by hemocytes [1–4]. The hemolymph of *Drosophila* is composed of three types of hemocytes: plasmatocytes (95%) (macrophages) have the capacity to remove foreign material by phagocytosis; crystal cells (5%) are involved in melanin synthesis during pathogen encapsulation [5] and lamellocyte, which are large flattened cells whose differentiation is induced in response to the immune system activation, i.e. the presence of foreign particles in the hemocoel.

The hemocytes of *Drosophila* are widely regarded as an excellent model for deciphering general innate immune mechanisms and DNA damage in animals [2,6–8]. *In vitro* and *in vivo* studies, no obvious toxicity of magnetic nanoparticles has been detected, but potential toxicity has been observed in blood and also activation of the immune systems [9].

Magnetite nanoparticles (Fe_3_O_4_NPs) are a common magnetic iron oxide that have an inverse spinal structure and the electrons can hop between 2+ and 3+ oxidation states of ions in the octahedral sites at room temperature, rendering magnetite an important class of half-metallic materials. With proper surface coating, these magnetic nanoparticles can be dispersed into suitable solvents, forming homogeneous suspensions called ferrofluids [10].

Due to the physicochemical properties of magnetite nanoparticles (Fe_3_O_4_NPs) and their application fields, biosafety information is insufficient and contradictory toxicity results have been reported due to different experimental conditions that would alter the effect of nanoparticles [11].

Chitosan (Ch) is the most abundant natural polysaccharide after cellulose and hemicelluloses. It is a non-toxic, biodegradable and biocompatible polysaccharide obtained from the deacetylation of chitin [12]. Chitosan provides nanoparticles with free amino and hydroxyl groups that enable the possibility to bind to a diversity of chemical groups and ions, leading to a number of applications such as protein and metal adsorption, guided drug and gene delivery, magnetic resonance imaging, tissue engineering and enzyme immobilization. Furthermore, this type of nanoparticle could be used in hyperthermia treatment for destroying malignant cells [13].

Chitosan in *Drosophila* has been well studied. *Drosophila* has been utilized for both production of chitosan [14] and as an *in vivo* model to investigate the transport and uptake of nanoparticles covered with chitosan in the larval digestive tract after oral administration [15].

In this study, we have observed *in vivo*, the cellular immune system activation in the hemolymph of *Drosophila* larvae by the effect of Ch-Fe_3_O_4_NPs exposure. To achieve these objectives, third instar larvae were exposed to two concentrations of Ch-Fe_3_O_4_NPs (500 and 1000 ppm) for 24 hours. Immune system activation was evaluated through hemolymph in terms of total number of hemocytes, apoptotic plasmatocytes, lamellocytes and DNA damage (comet assay). Additionally, the viability of larvae after the exposure to Ch-Fe_3_O_4_NPs was estimated.

## Materials and methods

### Synthesis and characterization of Ch-Fe_3_O_4_NPs

Ch-Fe_3_O_4_NPs were prepared by the protocol suggested by Gregorio *et al.* [16] with slight modifications. Transmission Electron Microscopy (TEM) micrographs were obtained using a FEI Tecnai G2 Spirit Twin at 80kV (Holland). Dynamic Light scattering (DLS) was conducted on diluted solutions previously filtered with a 220 nm PVDF filter membrane (Whatman, China), using the HORIBA LB-550 analyzer. The elemental analysis was obtained by EDX which was performed in the SEM chamber (Tescan Mira3) using a Bruker X-Flash 6|30 detector with a 123 eV resolution at Mn Kα. A sample was fixed in a stub previously covered with two layers of double coated carbon conductive tape and covering it with 20 nm of a conductive gold layer (99.99% purity) using a sputtering evaporator Quorum Q150R ES. The XRD measurement was carried out using an Empyrean diffractometer from PANalytical operating in a θ-2θ configuration (Bragg-Brentano geometry) and equipped with a Cu X-ray tube (Kα radiation λ = 1.54056 Å) operating at 40 kV and 40 mV.

### Exposure of *D. melanogaster* larvae to Ch-Fe_3_O_4_NPs

Third instar larvae of *Drosophila melanogaster* (Oregon R+ strain), were exposed to three treatments for 24 hours: 500 and 1000 ppm Ch-Fe_3_O_4_NPs, and a control without nanoparticles. Ch-Fe_3_O_4_NPs were supplied orally through the culture media. Fly cultures and larvae exposition took place in a 22°C incubator on a 12:12 light:dark cycle. The hemolymph of exposed larvae was extracted and analyzed to detect cell immune system activation through the total number of hemocytes, apoptotic plasmatocytes and lamelocytes as well as DNA damage (comet assay).

### Hemocytes counts

After the exposure, the hemolymph of thirty larvae was extracted and the hemocytes were stained with trypan blue 0,4 % (Santa Cruz Biotechnology). Three repetitions for each treatment were developed. Based on the morphology and color, normal hemocytes (transparent cells), apoptotic plasmatocytes (blue cells) and lamellocytes (large flat cells) were identified. The hemocytes were counted using a Neubauer chamber in a microscope ZEISS Imager A2 (40×/0.75).

The number of hemocytes (normal hemocytes, apoptotic plasmatocytes and lamellocytes) in larvae exposed to 500 ppm, 1000 ppm Ch-Fe_3_O_4_NPs and non-exposed was established.

Statistical differences between treatments were analyzed through a one way analysis of variance (ANOVA) in the SPSS software 23.0v (Windows Version 23.0. NY: IBM Corp. https://www-01.ibm.com/support/docview.wss?uid=swg21476197). The Bonferroni pos hoc test was developed to compare differences between nanoparticle treatments vs. the control test for each type of hemocytes. A probability less than 0.05 (p < 0.05) was considered statistically significant.

### Comet Assay

The comet assay in the alkaline version was developed in the hemocytes of larvae exposed to 500 ppm and 1000 ppm Ch-Fe_3_O_4_NPs and the control larvae according to the protocol described in Alaraby *et al.* [17]. The comets were visualized through a fluorescence microscopy Olympus DP72 using a 100×/0.17 lens.

A hundred hemocyte comets were observed for each treatment. Image captures and comet tail length were measured using the ImageJ software version 1.50e (https://imagej.net/Citing). The parameters used to estimate the DNA damage were: a) the percentage (%) of DNA in the comet tail and b) the tail length (μm).

The data was compared with an analysis of variance (ANOVA) test with the SPSS statistical software 23.0v. A probability of less than 0.05 (p < 0.05) was considered statistically significant. The Bonferroni post-hoc test was performed to compare the control versus the treatments exposed to Ch-Fe_3_O_4_NPs.

### Viability

A hundred of third instar larvae were exposed to each treatment (500 ppm and 1000 ppm, and control without Ch-Fe_3_O_4_NPs until adult eclosion. The progeny eclosioned from each treatment was counted. Non-eclosion after 8 days was counted as mortality. Three repetitions were performed for each treatment. The percentage of eclosioned flies in each treatment was compared using the SPSS statistical software 23.0v.

## Results

### Characterization of Ch-Fe_3_O_4_NPs

Figure 1 shows the TEM micrograph of Ch-Fe_3_O_4_NPs (1A) and the shape frequency histogram (1B). The average size of 155 measured nanoparticles by TEM was 11.0 ± 4.7 nm. This is compatible with the obtained DLS measurements: 9.2 ± 0.3 nm.

**Fig 1.**
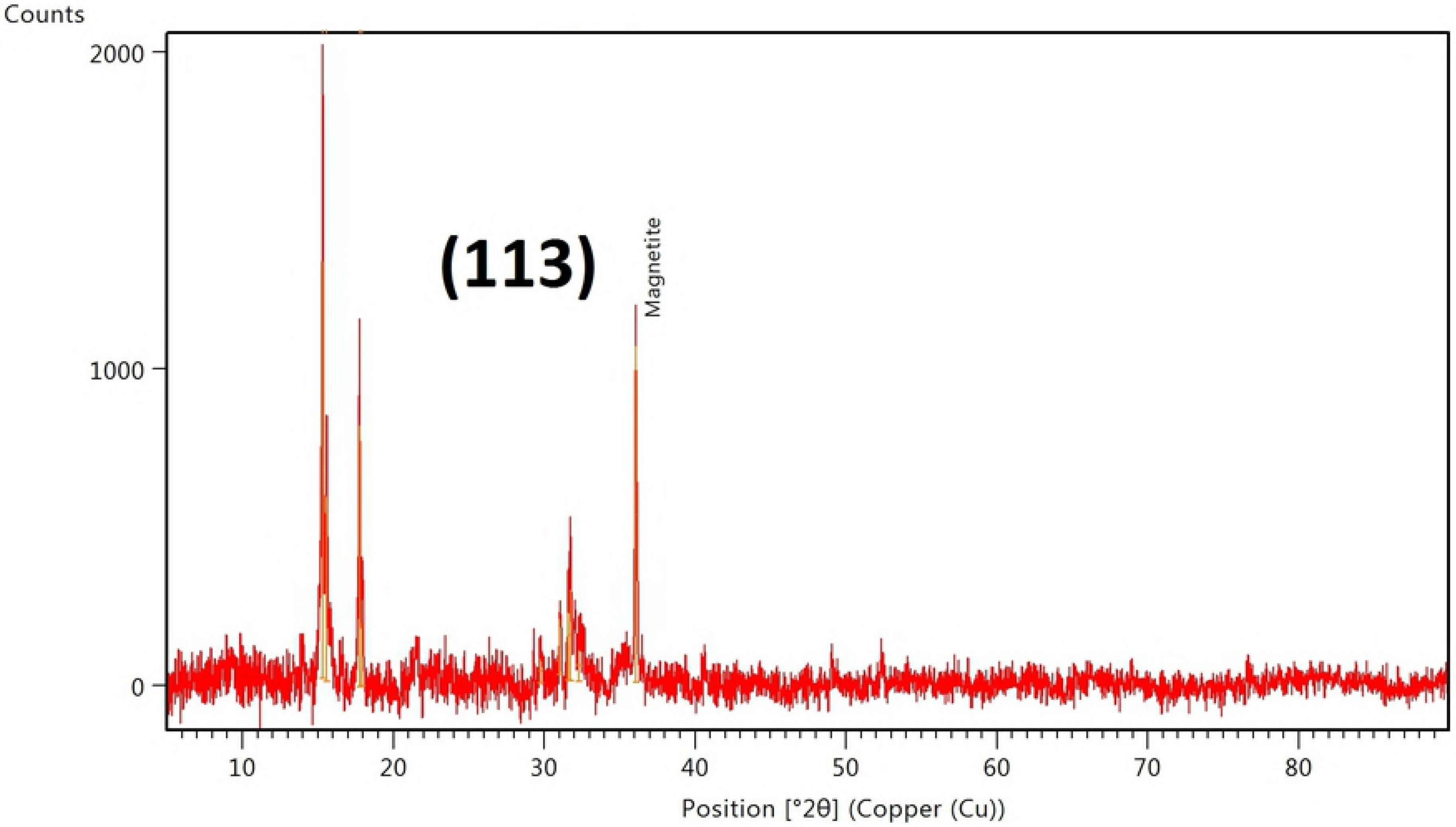
Micrographs of Ch-Fe_3_O_4_NPs. A) TEM and B) nanoparticles frequency histogram size.

The FEG-SEM micrograph (Fig 2) shows the chitosan recovering on Fe_3_O_4_NPs. The EDS measurements have been performed by considering C, N, O, Mg, S, Cl and Fe. In order to avoid biased determinations of the chemical compositions of the samples due to their inhomogeneity, we have averaged the spectra obtained from 25 points grid was averaged. The normalized weight average of each element and the standard deviation obtained by EDS analysis are listed in Table 1. We found the organic elements that comes from chitosan, C, N, and O. Chlorine comes from the inorganic salt precursors. Fe element comes from the magnetite nanoparticles. Traces of Mg and S come from the extraction process.

**Fig 2.**
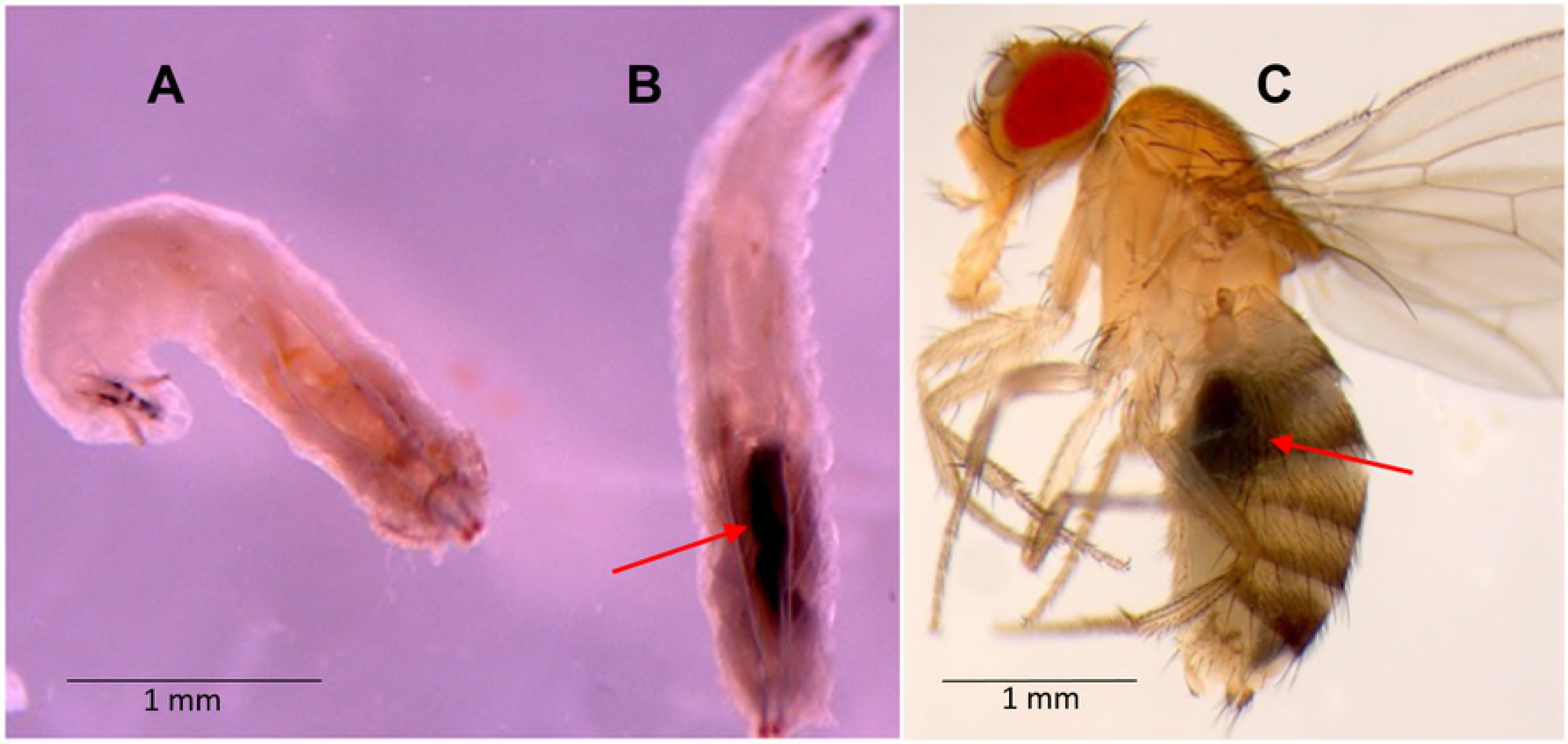
FEG-SEM micrograph of Ch-Fe_3_O_4_NPs.

**Table 1.**
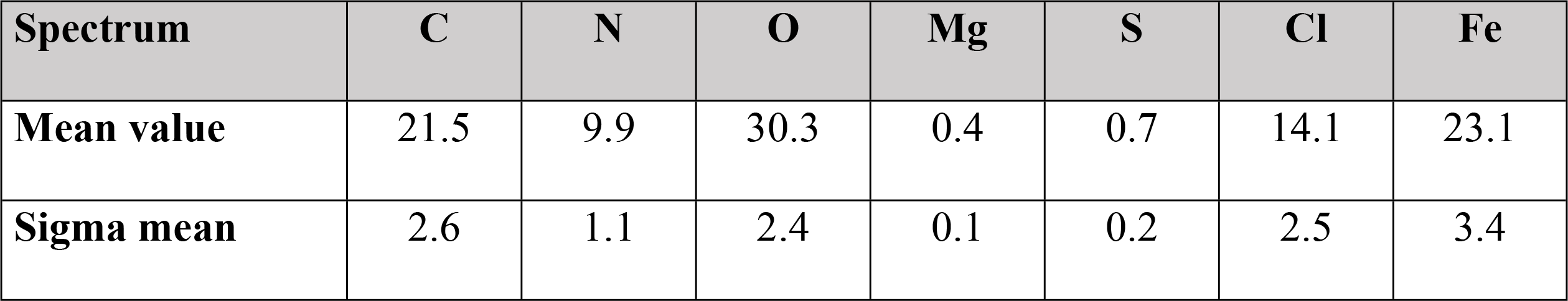
FEG-SEM EDX measurements of Ch-Fe_3_O_4_NPs in normalized wt.%

Samples of XRD were dried on a microscope slide at 40°C to avoid any organic degradation. Analysis XRD of the obtained average is the result of 6 different measurements from 5° to 90° (θ-2θ) angle. The Fe_3_O_4_NPs crystalline nature is confirmed from the XRD analysis (Fig 3). It is found that Bragg Reflection peaks at 36.06° which coincides with the cubic phase of Fe_3_O_4_ (ICSD: 96012). The lattice parameter and highest intensity plane (113) is well matched and agrees with other reported patterns [16]. Further peaks are observed around 15° and 20°. To our knowledge they correspond to impurities of the chitosan extract and its mix with the chemical compounds. Hematite or metal hydroxides were not identified, which confirms the complete formation of Fe_3_O_4_. The Debye Scherrer’s equation at the highest reflection peak (FWHM=0.168°) gives a 50 nm approximated size value for the Fe_3_O_4_ nanoparticles. This calculated value is higher than the TEM and DLS values likely due to the agglomeration of the organic extract.

**Fig 3.**
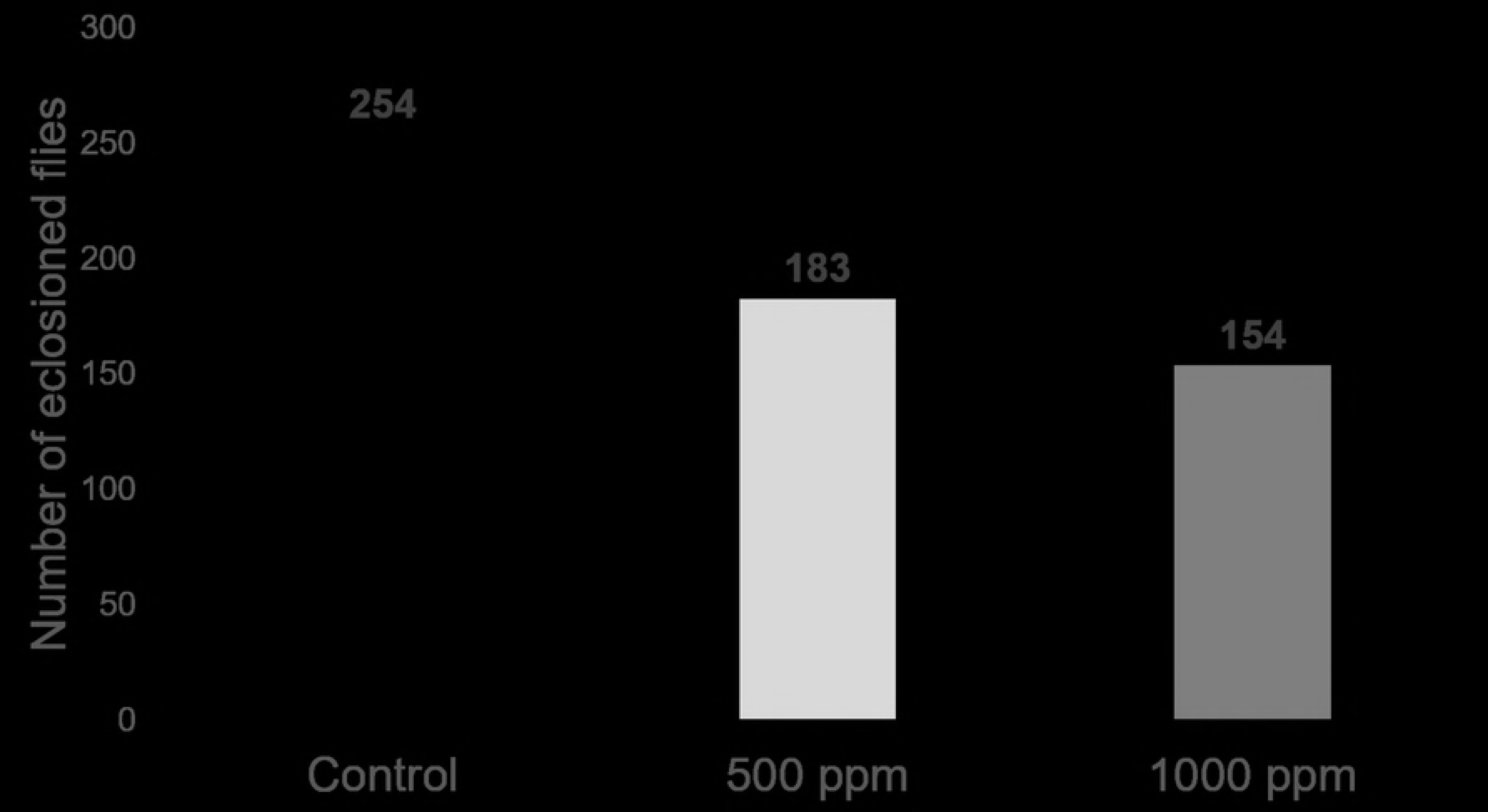
XRD pattern of the Ch-Fe_3_O_4_NPs.

### Hemocytes count

The changes in the total number of hemocytes and the presence of apoptotic or specialized cells were examined in order to detect the activation of immune system in the hemolymph of larvae exposed to Ch-Fe_3_O_4_NPs. Apoptotic hemocytes were identified by the blue coloration produced by entrance of trypan blue through the cell membrane. Lamellocytes were observed as large and irregular cells (Fig 4).

**Fig 4.**
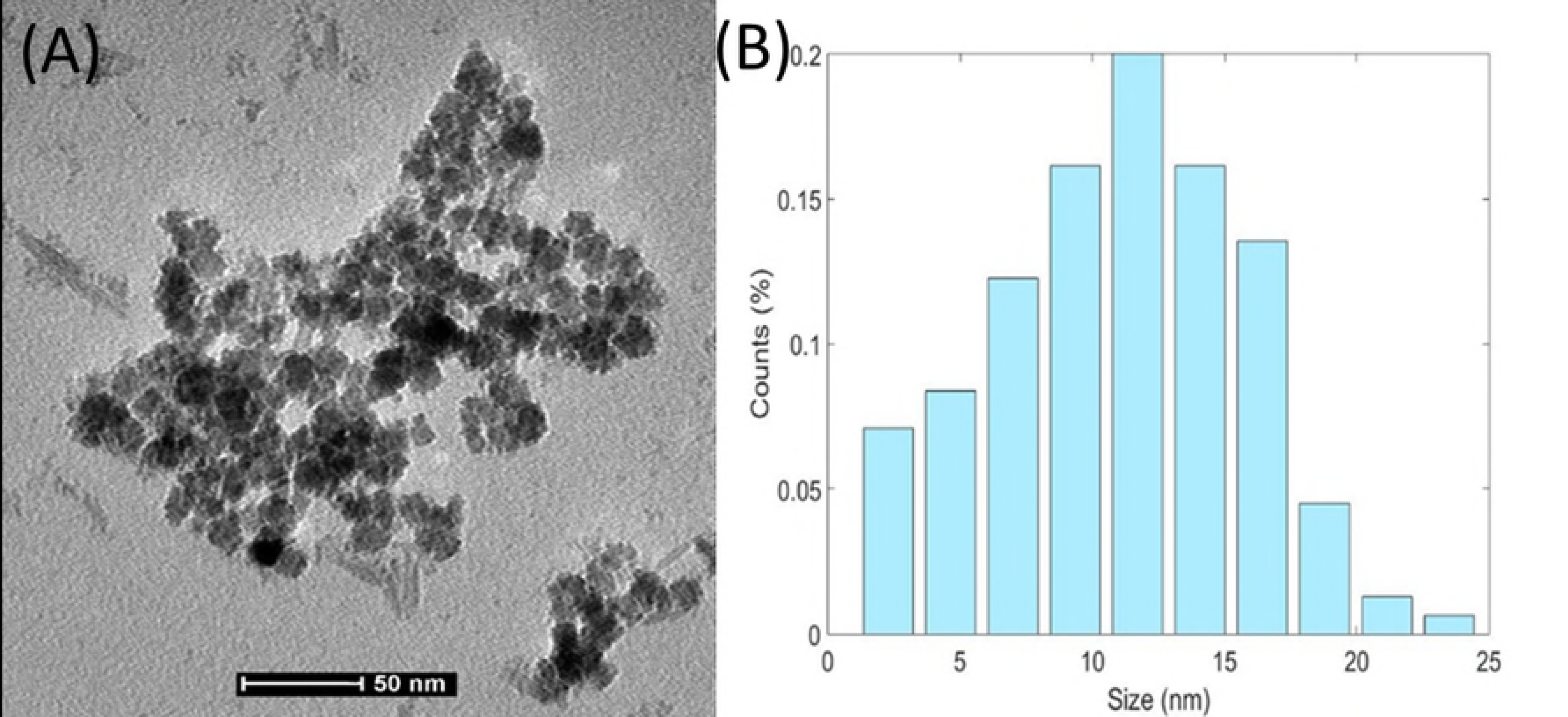
Hemolymph cells observed after nanoparticles exposure. A) normal plasmatocyte, B) normal lamellocyte, C) apoptotic plasmatocyte, D) apoptotic lamellocyte (40x/0.75).

The total number hemocytes increased in larvae exposed to 1000 ppm (mean: 411.33) but decreased in the larvae exposed to 500 ppm (mean: 201.67) compared with the control group (235.67). In the case of apoptotic plasmatocytes, the larvae exposed to 1000 ppm also showed an increase in the number of apoptotic plasmatocytes (mean: 54.33) compared with the 500 ppm (mean: 8.6) and control group (0.33). Lamellocytes were not present in the control larvae, but this type of cell was observed in the larvae exposed to 500 ppm (1.3) and 1000 ppm (13.3) (Fig 5).

**Fig 5.**
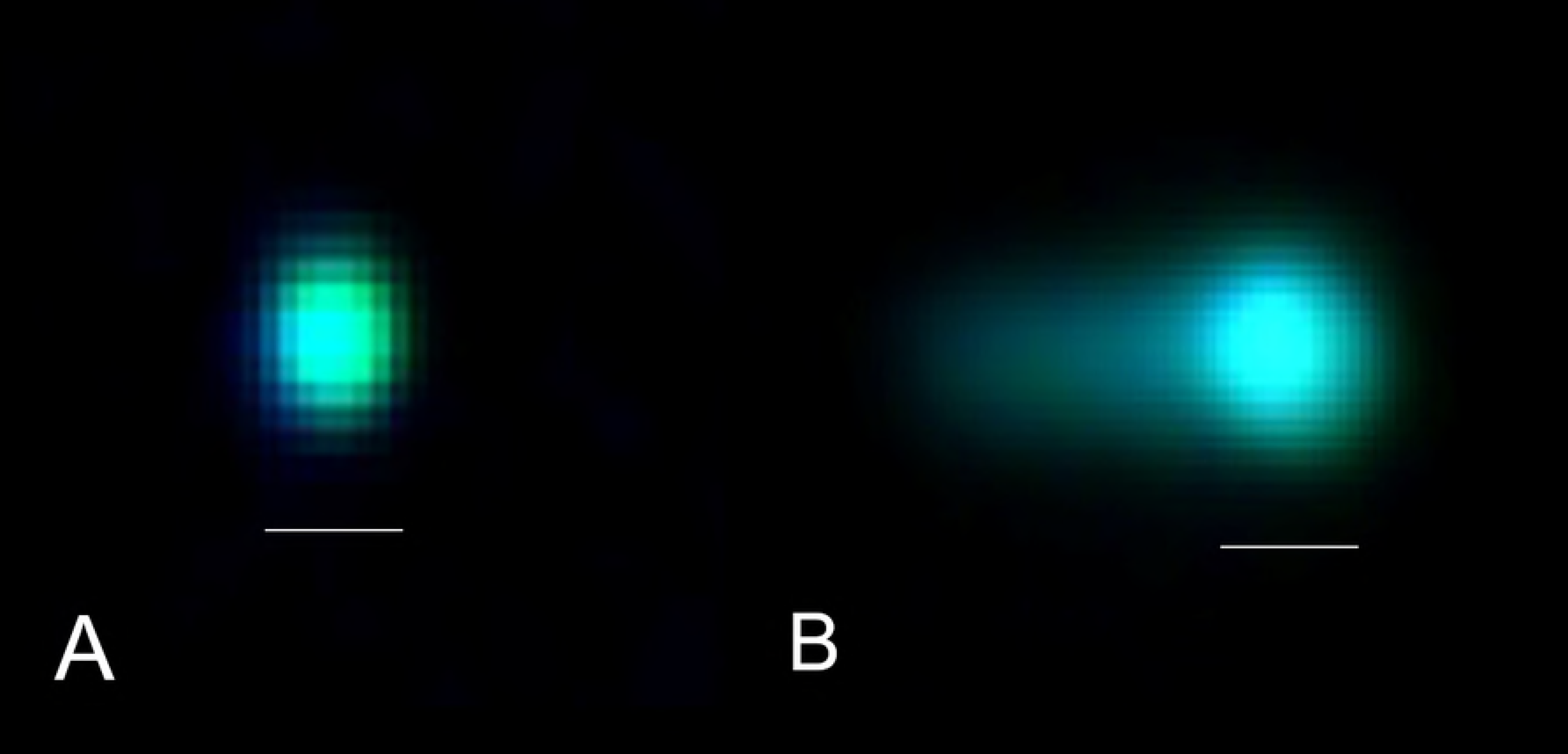
Hemocytes observed after nanoparticles exposure. Total number of hemocytes, apoptotic plasmatocytes and lamellocytes counted in each treatment.

For all counted cells (total number of hemocytes, apoptotic plasmatocytes and lamellocytes) (Table 2), statistical analysis shows little difference (p<0.05) between the larvae exposed to 500 ppm and the control test, but shows significant difference between the 1000 ppm treatment and the control test.

**Table 2.** Hemocytes counts per treatment.

| Treatment | Total hemocytes | Apoptotic plasmatocytes | Lamellocytes |
| --- | --- | --- | --- |
| Control | 707 | 1 | 0 |
| 500 ppm | 605 | 26 | 4 |
| 1000 ppm | 1234* | 163* | 40* |

\*Statistically significant $p < 0.05$

### Comet Assay

The comet assay was used to observe potential DNA damage in the hemocytes of larvae exposed to Ch-Fe_3_O_4_NPs. The DNA damage was detected by the presence of a comet tail in the cell nucleus. Comets without DNA damage and comets with high level of DNA damage were identified (Fig 6).

**Fig 6.**
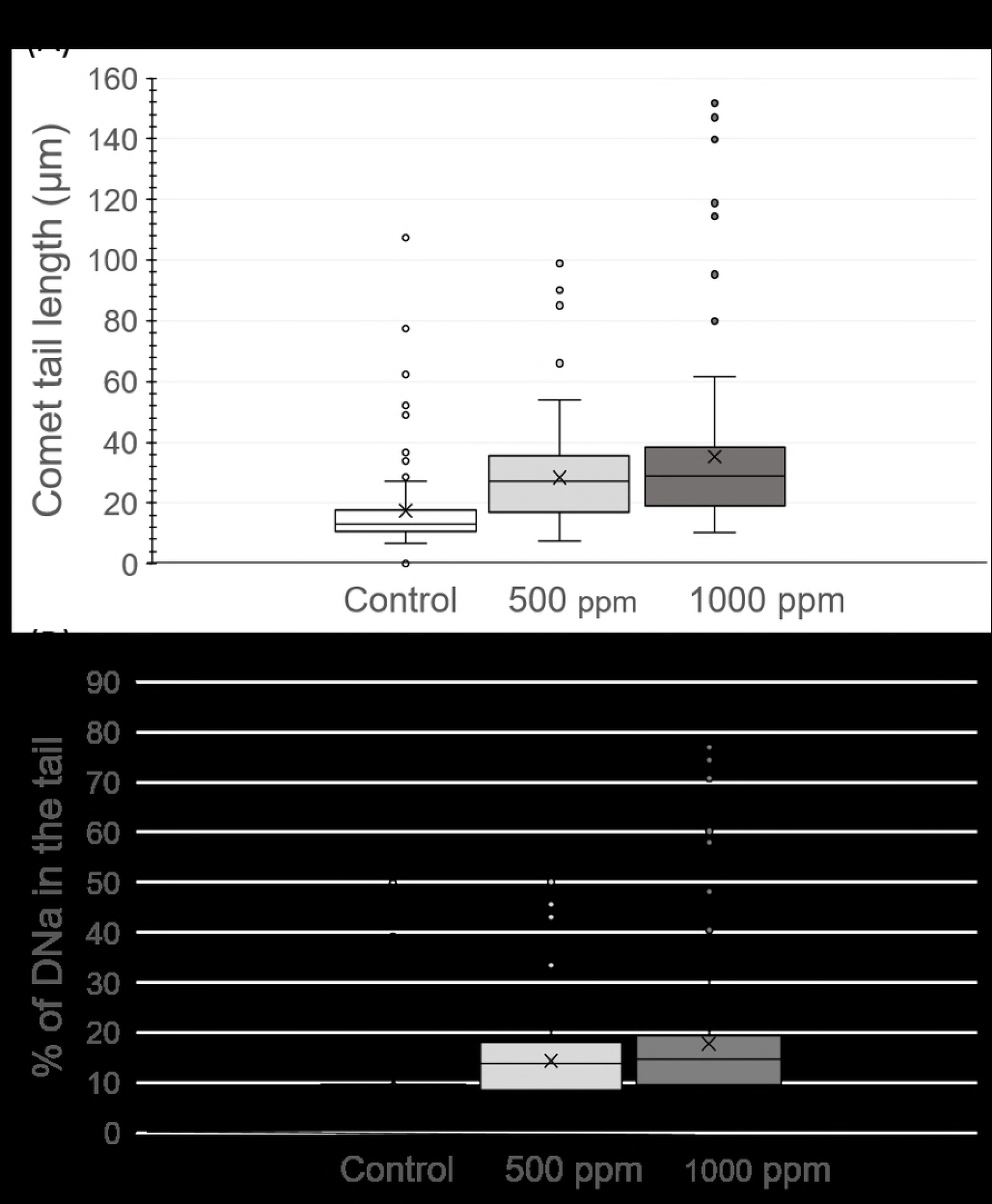
Nucleus observed in the comet assay. A) Hemocyte without comet tail, B) Hemocyte with comet tail and DNA damage. 400X (Bar = 25 μm)

DNA damage produced by exposure to each treatment in the hemocytes was estimated in function of the percentage of DNA (% of DNA) in the comet tail and the length of the comet tail. A direct association between Ch-Fe_3_O_4_NPs concentration and DNA damage was observed.

The level of DNA damage produced for each treatment was estimated in function of the % of DNA in the comet tail (Fig 7A) and the comet tail length (μm) (Fig 7B). The highest level of DNA damage was observed in the larvae exposed to 1000 ppm, followed by the larvae exposed to 500 ppm and finally the control larvae. However non-statistical differences were observed between the larvae exposed to 500 ppm and 1000 ppm (p<0.05). Therefore, both Ch-Fe_3_O_4_NPs concentrations are able to produce DNA damage in contrast with the control test (Table 3).

**Table 3.**
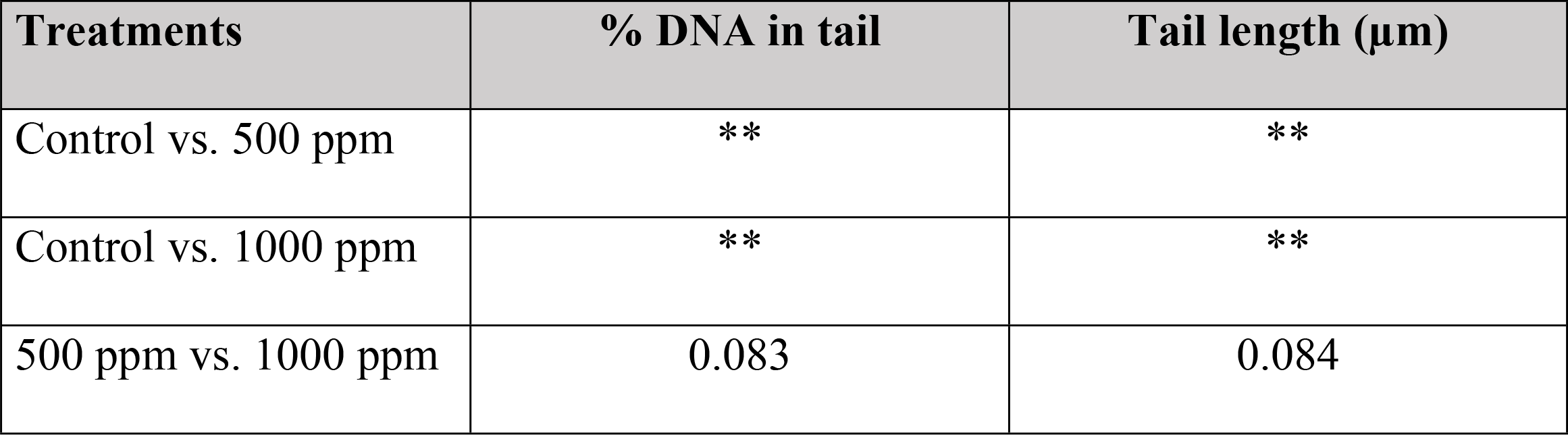
Statistical analysis of comet assay results.

**Fig 7.**
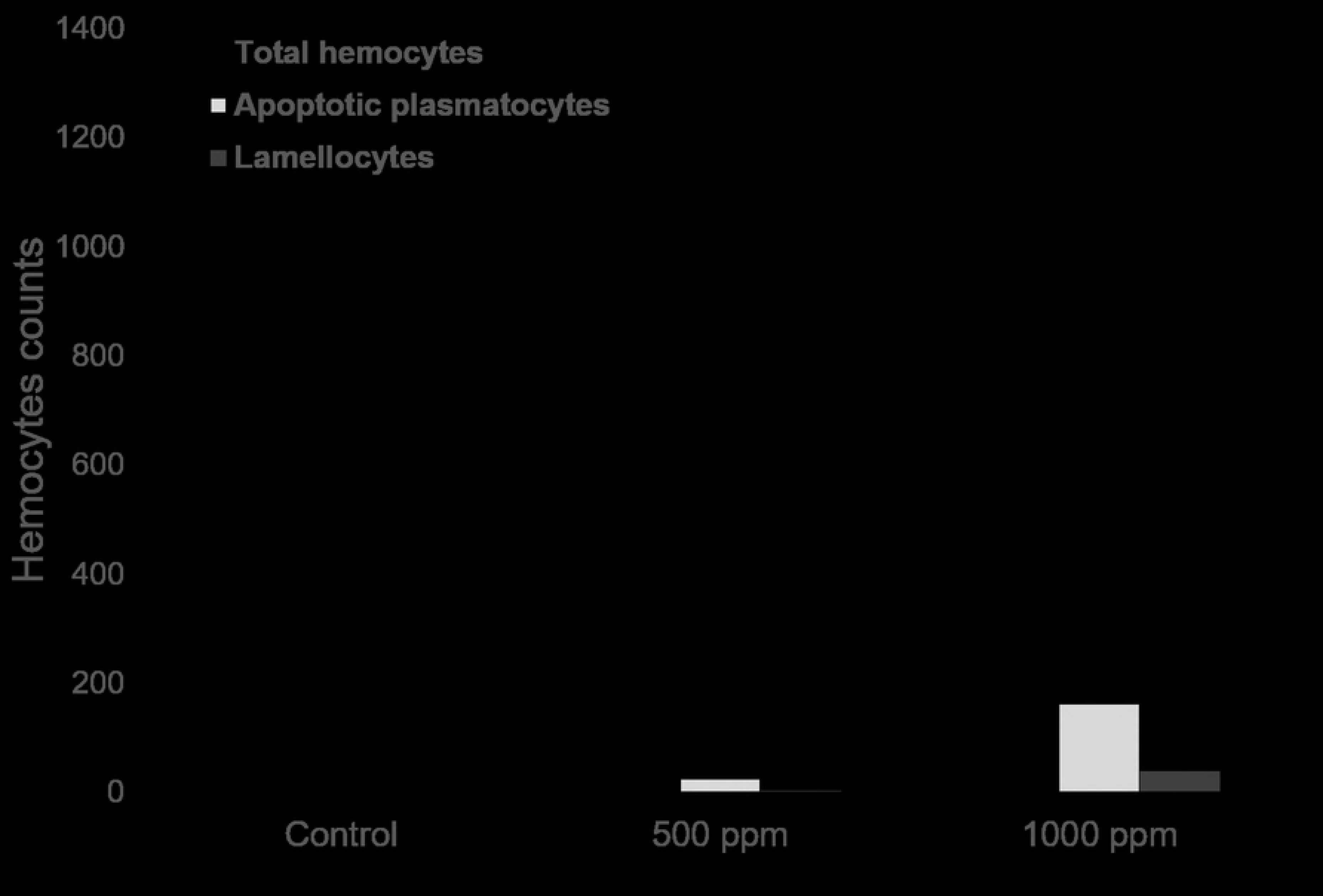
DNA damage observed by the comet test assay. Two parameters were used to estimate the DNA damage: A) % of DNA in the comet tail and B) comet tail length (μm)

### Viability of larvae

The viability of larvae was interpreted as the capacity of larvae to continue with the metamorphosis until the adult eclosion took place after the exposure to Ch-Fe_3_O_4_NPs. The parameter used to estimate the viability of larvae was the percentage of adults eclosioned after the exposure of the larvae to the Ch-Fe_3_O_4_NPs treatments. The lower percentage of viability was observed in the larvae exposed to 1000 ppm Ch-Fe_3_O_4_NPs (51.3%). A higher percentage was observed in the larvae exposed to 500 ppm (61.0%) and the highest percentage of viability was observed in the control test (84.0%). It is evident that mortality is directly associated to Ch-Fe_3_O_4_NPs concentration, therefore, exposure to high-dose concentration of Ch-Fe_3_O_4_NPs produce high mortality of larvae (Fig 8).

Significant statistical differences were observed in the viability between 1000 ppm and control treatment.

**Fig 8.**
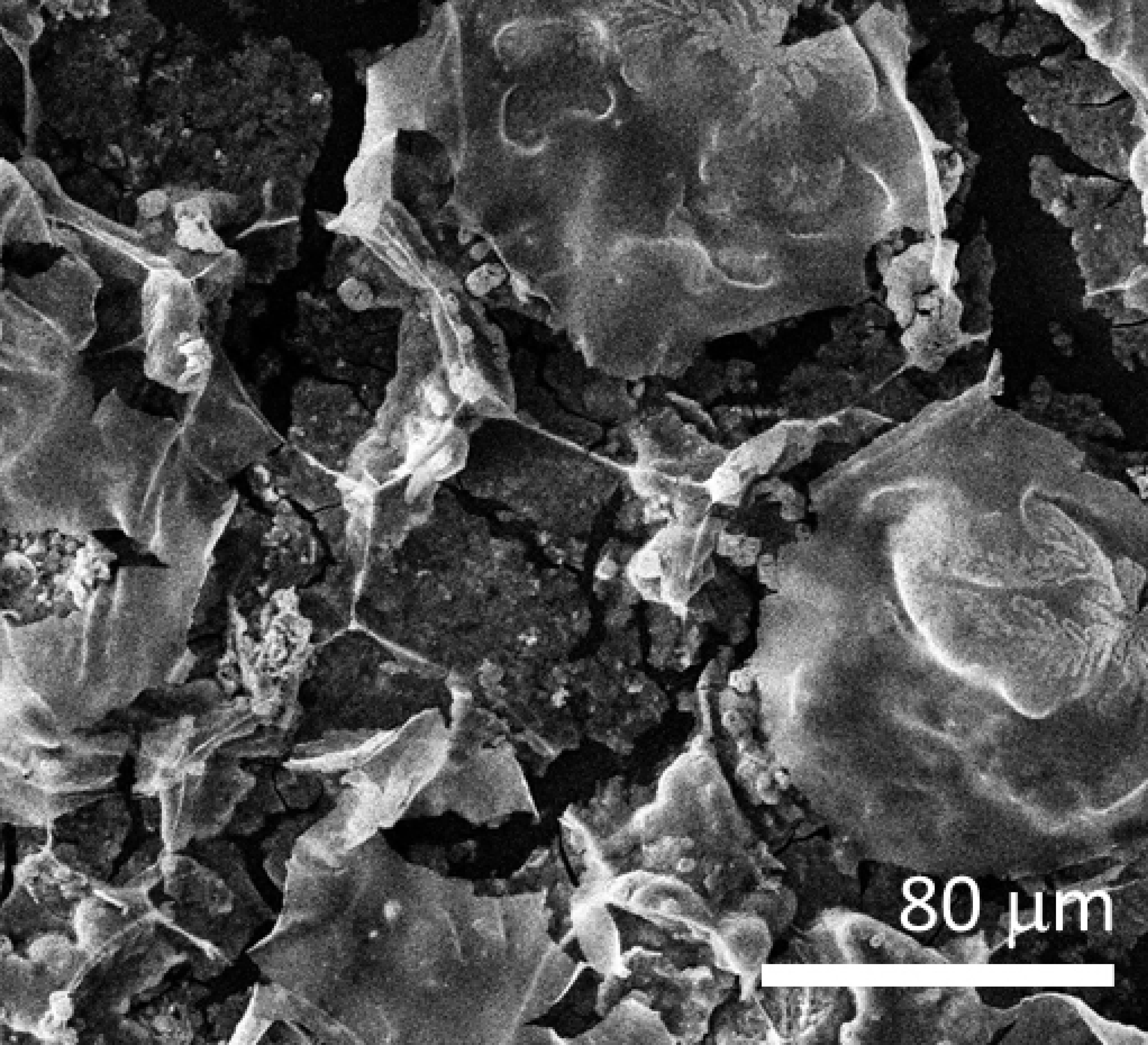
Viability of exposed larvaes. Adults eclosioned after exposure to each treatment.

Accumulation of Ch-Fe_3_O_4_NPs was observed in the midgut of the larvae after the exposure to the nanoparticles. This accumulation of Ch-Fe_3_O_4_NPs remains in the midgut of the eclosioned adults (Fig 9).

**Fig 9.**
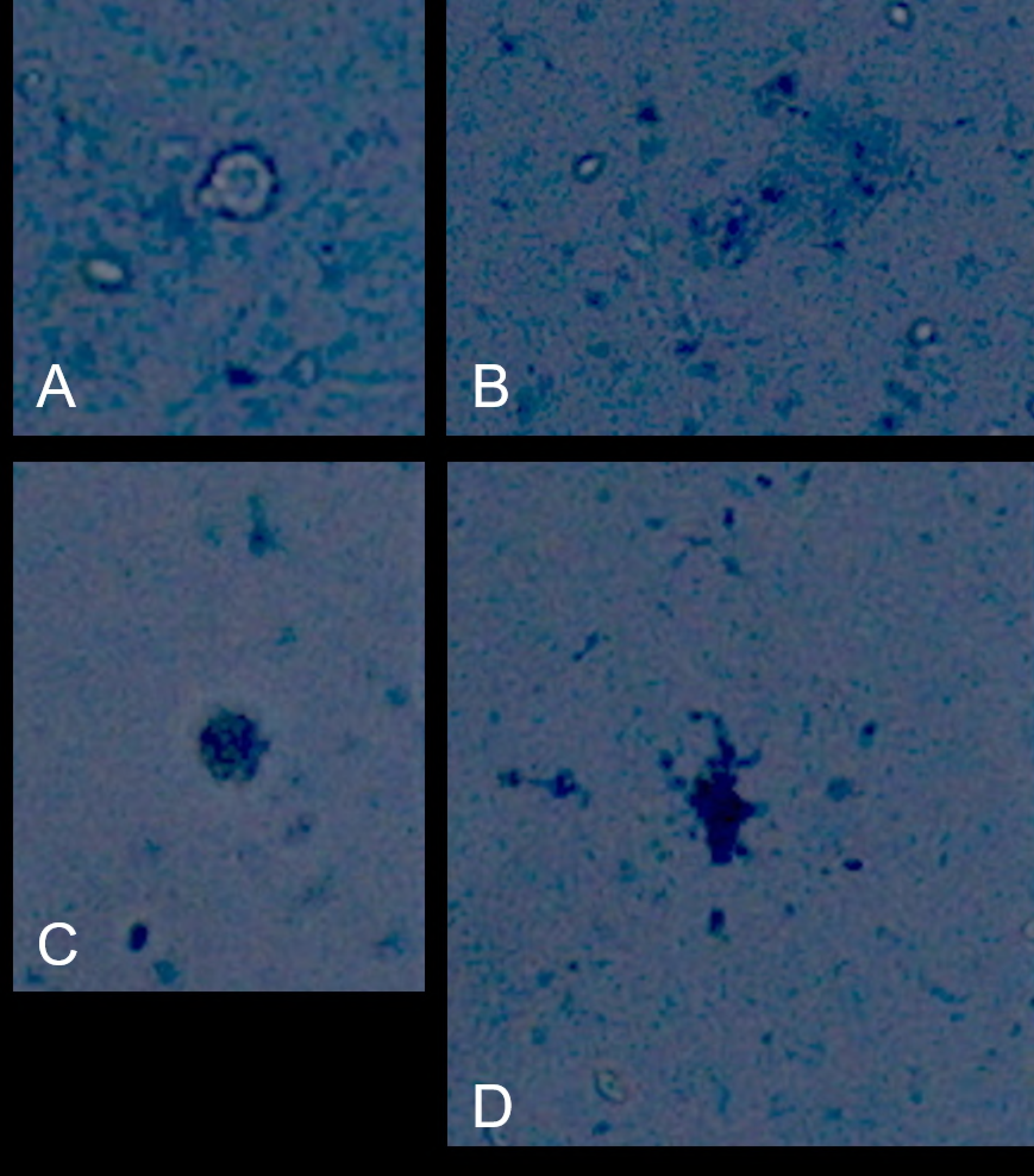
Accumulation of Ch-Fe_3_O_4_NPs in the midgut of third instar larvae as an adult. A) larva not exposed (control), B) larvae exposed to 1000 ppm, C) adult exposed to 1000 ppm Ch-Fe_3_O_4_NPs (red arrows show accumulation of Ch-Fe_3_O_4_NPs).

## Discussion

In *Drosophila* the cellular immune response starst in the hemolymph through the hemocytes, and the circulating immune surveillance cells play a central role in the immune response. In this response each type of hemocyte has a function. When an invading organism or particle is recognized as foreign, circulating plasmatocytes may remove it by phagocytosis and lamellocytes can complete the process by encapsulation when the particles are too large to undergo phagocytosis [18].

In this study changes were observed in the density of cell hemolymph composition in *Drosophila* larvae exposed for 24 hours to Ch-Fe_3_O_4_NPs including the increment of the number of plasmatocytes, the emergence of lamellocytes and the presence of apoptotic plasmatocytes. Additionally, DNA damage and high mortality of larvae exposed to Ch-Fe_3_O_4_NPs were observed.

The increment of plasmatocytes density and the emergence of lamellocytes were observed in larvae exposed to 500 ppm and 1000 ppm. However the effect of 1000 ppm concentration is toxic, while the exposure to 500 ppm is less nocive for the hemolymph cells. This observation could be explained due to the presence of *Drosophila* hemocytes: circulating (in hemolymph) and sessile hemocytes (in the body wall) which have different functions during the immune response. The circulating plasmatocytes in the larvae are originated through prohemocytes (embrionic macrophages); this differentiation occurs during the normal development of the larvae. However, when an event like the presence of pathogens, parasites or foreign particles are detected, the cellular immune response is activated [19], and the sessile hemocytes detach from the epithelium and enter the circulating hemolymph triggering the differentiation of plasmatocytes or lamellocytes, increasing the number of circulating hemocytes (plasmatocytes and lamellocytes). The emergence of lamellocytes could be originated by both prohemocytes differentiation and also by plasmatocytes differentiation. The plasmatocyte differentiation to originated lamellocytes is triggered by immune induction [18,20,21]. Therefore, the increase of the number of circulating plasmatocytes and the emergence of lamellocytes are evident signals of cellular immune response activation.

Larvae exposed to 1000 ppm of Ch-Fe_3_O_4_NPs showed the highest number of emergent lamellocytes (163 cells), the highest number of apoptotic hemocytes (40 cells) and the highest level of DNA damage. This suggests that high concentrations of nanoparticles could produce toxic effects in the hemocytes. This dose-concentration effect of Ch-Fe_3_O_4_NPs is supported by the statistical analysis that shows no difference between the 500 ppm treatment and the control but high significant difference between the 1000 ppm treatment and the control. The toxic effect of Ch-Fe_3_O_4_NPs at 1000 ppm concentration was demonstrated in this study. The dose concentration and the exposure time are important factors that influence the level of toxicity in hemocytes [22]. Also, the type of hemocyte is related to the toxic effect due to different structural characteristic and function of each hemolymph cells [23].

Another signal of immune system activation is the presence of apoptotic cells which is a signal of hemocyte damage and hemocyte death due to cell membrane damage, and plays a key role in immune response by eliminating cells subjected to various stress factors [24]. In this study, the high number of apoptotic cells were observed in larvae exposed to 1000 ppm Ch-Fe_3_O_4_NPs. This damage was observed by the increase of blue stained hematocytes in larvae exposed to Ch-Fe_3_O_4_NPs.

Also, the toxic effect of Ch-Fe_3_O_4_NPs on the DNA was demonstrated through the comet assay which allows for associating the percentage of DNA in the tail and the length of comet tail with the level of DNA damage. Exposure to 1000 ppm Ch-Fe_3_O_4_NPs produced the highest percentage of DNA in the comet and the highest length of comet tail. Comet assay provides a sensitive way to detect the effects of Ch-Fe_3_O_4_NPs by means of measuring DNA strand breaks [25,26]; this will allow identification of the possible mode of action of nanoparticles at the molecular level.

In addition, the effect of Ch-Fe_3_O_4_NPs on the viability of larvae was evidently toxic, producing up to 50% of mortality in larvae exposed to 1000 ppm Ch-Fe_3_O_4_NPs, while the non-exposed larvae presented only up to 16% of mortality. The low viability is associated to the Ch-Fe_3_O_4_NPs exposure; however, the physiological mechanisms should be analyzed in future research.

Accumulation of Ch-Fe_3_O_4_NPs in the midgut of exposed larvae was observed; this accumulation was transferred to the adult during the metamorphosis. This event could be associated to the immune response through cellular events such as phagocytosis and humoral events that include lysis and melanization [27]. Another explanation could be related to the low capacity of larvae to excrete the nanoparticles. The effect of this accumulation has not been analyzed in this study. However new studies should be conducted to establish if agglomeration of Ch-Fe_3_O_4_NPs in the digestive tract are produced by plasmatocytes through phagocytosis, and additionally accumulation of nanoparticles should be estimated.

## Conclusion

Activation of cellular immune response was observed in the hemolymph of *Drosophila* larvae through the increment of hemocytes density, the emergence of lamellocytes and the presence of apoptotic hemocytes after the exposure to Ch-Fe_3_O_4_NPs. In addition, DNA damage detected in hemocytes by the comet assay, and the low viability of larvae is directly associated to the dose concentration Ch-Fe_3_O_4_NPs. The toxic effect of nanoparticles is higher in larvae exposed to 1000 ppm concentration, while 500 ppm could have toxic risks but have not been detected in this study.

## References

1. Gillespie, Jeremy P.; Kanost MR. Biological Mediators of Insect Immunity. Annu Rev Entomol. 1997;42:611–43.

2. Irving P, Ubeda JM, Doucet D, Troxler L, Lagueux M, Zachary D, et al. New insights into Drosophila larval haemocyte functions through genome-wide analysis. Cell Microbiol. 2005;7:335–50.

3. Lackie AM. Haemocyte Behaviour. Adv. In Insect Phys. 1988.

4. Strand M, Pech L. Immunological Basis for Compatibility in parasitoid-host relationships. Annu Rev Entomol. 1995;40:31–56.

5. Söderhäll K, Cerenius L. Role of the prophenoloxidase-activating system in invertebrate immunity. Curr Opin Immunol. 1998;10:23–8.

6. Carmona ER, Inostroza-Blancheteau C, Obando V, Rubio L, Marcos R. Genotoxicity of copper oxide nanoparticles in Drosophila melanogaster. Mutat Res - Genet Toxicol Environ Mutagen [Internet]. Elsevier Ltd; 2015;791:1–11. Available from: http://dx.doi.org/10.1016/j.mrgentox.2015.07.006

7. Lemaitre B, Hoffmann J. The Host Defense of Drosophila melanogaster. Annu Rev Immunol [Internet]. 2007;25:697–743. Available from: http://www.annualreviews.org/doi/10.1146/annurev.immunol.25.022106.141615

8. Cherry S, Silverman N. Host-pathogen interactions in drosophila: New tricks from an old friend. Nat Immunol. 2006;7:911–7.

9. Elsabahy M, Wooley KL. Data Mining as a Guide for the Construction of Cross-Linked Nanoparticles with Low Immunotoxicity via Control of Polymer Chemistry and Supramolecular Assembly. Acc Chem Res. 2015;48:1620–30.

10. Odenbach S. Ferrofluids and their applications. MRS Bull. 2013;38:921–4.

11. Lei C, Sun Y, Tsang DCW, Lin D. Environmental transformations and ecological effects of iron-based nanoparticles. Environ Pollut [Internet]. Elsevier Ltd; 2017;232:10–30. Available from: http://linkinghub.elsevier.com/retrieve/pii/S0269749117322601

12. Zhu A, Yuan L, Liao T. Suspension of Fe3O4 nanoparticles stabilized by chitosan and o-carboxymethylchitosan. Int J Pharm. 2008;350:361–8.

13. Haldorai Y, Kharismadewi D, Tuma D, Shim JJ. Properties of chitosan/magnetite nanoparticles composites for efficient dye adsorption and antibacterial agent. Korean J Chem Eng. 2015;32:1688–93.

14. Kaya M, Akyuz B, Bulut E, Sargin I, Eroglu F, Tan G. Chitosan nanofiber production from Drosophila by electrospinning. Int J Biol Macromol. 2016;92:49–55.

15. Nimal TR, Baranwal G, Bavya MC, Biswas R, Jayakumar R. Anti-staphylococcal Activity of Injectable Nano Tigecycline/Chitosan-PRP Composite Hydrogel Using Drosophila melanogaster Model for Infectious Wounds. ACS Appl Mater Interfaces. 2016;8:22074–83.

16. Gregorio-Jauregui KM, Pineda MG, Rivera-Salinas JE, Hurtado G, Saade H, Martinez JL, et al. One-step method for preparation of magnetic nanoparticles coated with chitosan. J Nanomater. 2012;2012.

17. Alaraby M, Annangi B, Hernández A, Creus A, Marcos R. A comprehensive study of the harmful effects of ZnO nanoparticles using Drosophila melanogaster as an in vivo model. Hazard Mater. 2015;296:166–74.

18. Honti V, Csordás G, Kurucz É, Márkus R, Andó I. The cell-mediated immunity of Drosophila melanogaster: Hemocyte lineages, immune compartments, microanatomy and regulation. Dev Comp Immunol [Internet]. Elsevier Ltd; 2014;42:47–56. Available from: http://dx.doi.org/10.1016/j.dci.2013.06.005

19. Zettervall C-J, Anderl I, Williams MJ, Palmer R, Kurucz E, Ando I, et al. A directed screen for genes involved in Drosophila blood cell activation. Proc Natl Acad Sci [Internet]. 2004;101:14192–7. Available from: http://www.pnas.org/cgi/doi/10.1073/pnas.0403789101

20. Honti V, Csordás G, Márkus R, Kurucz É, Jankovics F, Andó I. Cell lineage tracing reveals the plasticity of the hemocyte lineages and of the hematopoietic compartments in Drosophila melanogaster. Mol Immunol. 2010;47:1997–2004.

21. Stofanko M, Kwon SY, Badenhorst P. Lineage tracing of lamellocytes demonstrates Drosophila macrophage plasticity. PLoS One. 2010;5.

22. Xing R, Li K Le, Zhou YF, Su YY, Yan SQ, Zhang KL, et al. Impact of fluorescent silicon nanoparticles on circulating hemolymph and hematopoiesis in an invertebrate model organism. Chemosphere [Internet]. Elsevier Ltd; 2016;159:628–37. Available from: http://dx.doi.org/10.1016/j.chemosphere.2016.06.057

23. Li K-L, Zhang Y-H, Xing R, Zhou Y-F, Chen X-D, Wang H, et al. Different toxicity of cadmium telluride, silicon, and carbon nanomaterials against hemocytes in silkworm, Bombyx mori. RSC Adv [Internet]. Royal Society of Chemistry; 2017;7:50317–27. Available from: http://xlink.rsc.org/?DOI=C7RA09622D

24. Gervais O, Renault T, Arzul I. Induction of apoptosis by UV in the flat oyster, Ostrea edulis. Fish Shellfish Immunol [Internet]. 2015;46:232–42. Available from: http://www.sciencedirect.com/science/article/pii/S1050464815300255%5Cn http://ac.els-cdn.com/S1050464815300255/1-s2.0-S1050464815300255-main.pdf?_tid=a1466e7a-36d5-11e5-96ec-00000aacb35f&acdnat=1438272866_a1221c9be17ba9a7e2935794e93dea25

25. Collins a R. The comet assay for DNA damage and repair: principles, applications, and limitations. Mol Biotechnol [Internet]. 2004;26:249–61. Available from: http://www.ncbi.nlm.nih.gov/pubmed/15004294

26. Augustyniak M, Gladysz M, Dziewiecka M. The Comet assay in insects-Status, prospects and benefits for science. Mutat Res - Rev Mutat Res. 2016;767:67–76.

27. Hillyer JF. Insect immunology and hematopoiesis. Dev Comp Immunol [Internet]. Elsevier Ltd; 2016;58:102–18. Available from: http://dx.doi.org/10.1016/j.dci.2015.12.006

